# Population genomics reveals mechanisms and dynamics of *de novo* proto-gene emergence in *Drosophila melanogaster*

**DOI:** 10.1101/2022.11.19.517182

**Authors:** Anna Grandchamp, Lucas Kühl, Marie Lebherz, Kathrin Brüggemann, John Parsch, Erich Bornberg-Bauer

## Abstract

Novel genes are essential for evolutionary innovations and differ substantially even between closely related species. Recently, multiple studies across many taxa have suggested that some novel genes arise *de novo*, i.e. from previously non-coding DNA. In order to characterise the underlying mutations that allowed *de novo* gene emergence and their order of occurrence, homologous regions must be detected within non-coding sequences in closely related sister genomes. So far, most studies do not detect non-coding homologs of *de novo* genes due to inconsistent data and long evolutionary distances separating genomes. Here we overcome these issues by searching for proto-genes, the not-yet fixed precursors of *de novo* genes that emerged within a single species. We sequenced and assembled genomes with long-read technology and the corresponding transcriptomes from inbred lines of *Drosophila melanogaster*, derived from seven geographically diverse populations. We found line-specific proto-genes in abundance but few proto-genes shared by lines, suggesting a rapid turnover. Gain and loss of transcription is more frequent than the creation of Open Reading Frames (ORFs), e.g. by forming new START- and STOP-codons. Consequently, the gain of ORFs becomes rate limiting and is frequently the initial step in proto-gene emergence. Furthermore, Transposable Elements (TEs) are major drivers for intra genomic duplications of proto-genes, yet TE insertions are less important for the emergence of proto-genes. However, highly mutable genomic regions around TEs provide new features that enable gene birth. In conclusion, proto-genes have a high birth-death rate, are rapidly purged, but surviving proto-genes spread neutrally through populations and within genomes.

## Introduction

*De novo* gene origination is a recently recognised process that describes the emergence of new genes from previously non-coding sequences via a series of mutational events (Levy, 2019; Kaessmann, 2010; Schlötterer, 2015). It has been conjectured that over very short evolutionary time scales, most novel genes originate *de novo* but are then rapidly lost (Rödelsperger et al., 2019; Schmitz et al., 2018; Lange et al., 2021). Most probably, newly emerged *de novo* genes fail to establish a sufficient advantage to the host organism, and thus are easily purged by drift or negative selection. Over longer time scales, most observed novel genes stem from duplication events. Proteins derived from duplicated genes are more likely to have already approved structures and functions which do not negatively interfere with the long established cellular network (O’Toole et al., 2018), and gain new functional roles, through mechanisms such as sub- or neo-functionalisation (Innan and Kondrashov, 2010; Rastogi and Liberles, 2005; Konrad et al., 2011).

The emergence of functional proteins from *de novo* genes is difficult to rationalise with our current understanding of molecular genetics and evolution. Although believed to be highly unlikely until recently, *de novo* gene birth has by now been described in many species, including several fungi, plants, insects, mammals and fishes (Li et al., 2016; Ruiz-Orera et al., 2015; Wu and Knudson, 2018; Zhao et al., 2014). Even more intriguingly, some genes that emerge *de novo* not only provide new functions, but have even become essential to the species in which they emerged. For example, the human-specific *de novo* gene ESRG is required for the maintenance of pluripotency in human naive stem cells (Wang et al., 2014). MDF1, a *de novo* gene found only in *S. cerevisiae*, has a function in suppressing sexual reproduction (Li et al., 2014). In humans, *de novo* genes have been shown to be utilized for brain development (Li et al., 2010). Overall, many of the predicted *de novo* genes in metazoa seem to be associated with either developmental processes of the neuronal system or with reproduction. Both of these processes are well known for their fast genetic turnover and adaptation (Kaessmann, 2010; Maze et al., 2015; Cutter et al., 2019). Several predicted *de novo* genes in the Drosophila genus were shown to have become essential for male fertility (Rivard et al., 2021; Gubala et al., 2017).

To ascertain that a candidate *de novo* gene did not emerge via a duplication or transposition event, the gene must not show homology to any other gene in the species in which it emerged and to any other gene in an outgroup species of the taxonomic group under study. However, as duplicated genes are more likely to evolve faster than single-copy genes (O’Toole et al., 2018), a duplicated gene could also lack a detectable orthology relationship. Most earlier studies of *de novo* genes relied on the comparative ge-nomics of species that diverged several tens of millions of years (my) ago (Schmitz et al., 2018; Dowling et al., 2020; Neme and Tautz, 2013; Begun et al., 2006; Cai et al., 2008; McLysaght and Guerzoni, 2015). This led to considerable discussions regarding the reliability of their *de novo* status (Moyers and Zhang, 2016; Domazet-Lošo et al., 2017; Weisman et al., 2020).

The only way to conclusively demonstrate that a gene emerged from scratch is to locate the corresponding homologous non-coding sequence in the closest outgroup species and study the putative mutations that transformed a non-coding sequence into a coding one. For a proto-gene to emerge, at least two events must occur: the emergence of a new ORF, and the gain of transcription. Nonetheless, genomes and synteny between two closely related species can be extensively reshuffled during evolution. The extensive divergence of non-coding regions due to lack of purifying selection implies that, the deeper the phylogeny becomes, the less likely the non-coding regions that are homologous to a predicted *de novo* gene can be identified. More sensitive phylogenetic procedures that rigorously refine the first-trawl-results from phylostratigraphic analyses have been suggested and implemented (McLysaght and Guerzoni, 2015; Wu and Knudson, 2018; Heames et al., 2020). As a key feature, they employ gene syntenies to allow for highly sensitive matches (i.e those with little sequence similarity) if *de novo* genes appear between the same gene pairs in two or more genomes. Using synteny works well in mammals (Jebb et al., 2020; Vakirlis et al., 2020) or in vertebrates in general but gene order is less conserved in taxa such as insects (Zdobnov and Bork, 2007).

An improved strategy to study the emergence of *de novo* genes is to study genomes of high quality covering a very narrow time scale. Using several rice genomes, Zhang et al. (2019) managed to pinpoint precisely the enabling mutations allowing emergence of *de novo* genes in closely related rice species. A recent state-of-the-art study used six nematode populations and confirmed the rapid turnover of *de novo* genes compared to duplicates (Prabh and Rödelsperger, 2022). Several other studies included fewer species but focused on even shorter evolutionary time scales by using population data. By analysing population data from wild fish populations, it has been corroborated frequent random emergence (and rapid loss) of transcripts and that (few) surviving transcripts gain stable and broader expressions, presumably via fast epigenetic control (Schmitz et al., 2020). Zhao et al. (2014) identified 106 *de novo* genes specifically expressed in the testis of laboratory strains of *D. melanogaster*. Furthermore, Moyers and Zhang (2016) demonstrated that 13.9% of *D. melanogaster* genes did not have a detectable ortholog in any outgroup species, and Heames et al. (2020) reported that 26.8% of the transcribed ORFs of *D. melanogaster* originated from orphan genes. Despite this progress, all of these studies are hampered by either data disparity, the use of discontinuous evolutionary time scales, a lack of precisely annotated genomes for inter- and intra-specific comparison, or several of these issues. For these reasons, the full extent of *de novo* genes in a species, the mutations enabling their emergence, and their prevalence in natural populations, remain unknown.

In the present study, we overcome several of the limitations of earlier studies by using in-bred lines of *D. melanogaster* from various geographic locations to generate line specific genomes and transcriptomes. In doing so, we avoid issues of intra-specific allelic variation and provide an early snapshot of the origin of *de novo* genes. The small size of the genomes and the availability of a well annotated reference genome allows us to localize most *de novo* genes at the chromosomal level and to follow mutations and structural variations associated with the very early stages of *de novo* gene emergence. By using long-read sequence data (Nanopore) and a common annotation strategy across all genomes we also avoid recently recognised issues arising from the use of disparate methods (Weisman et al., 2022). Overall, we detect al large number of *de novo* genes in all lines, demonstrate their fast turnover and, by using homology and synteny approaches, detect and characterise the enabling mutations underlying their emergence. Since most of these *de novo* genes have not yet become fixed within a species and are neither stably transcribed or translated across species boundaries, we refer to them as *proto-genes*, following the terminology proposed by Carvunis et al. (2012).

## Results

### Genome sequencing, assembly and annotation

We studied proto-gene emergence in 7 inbred lines of *D. melanogaster*, including 6 from derived European populations and one from the ancestral species range in Zambia (Figure 1a; supplemental Table S1.xls), using Nanopore long-read sequencing. We obtained 9-21 Gb of raw DNA sequence per line (Genbank accession code, in progress), resulting in 66-158 X coverage (Table 1). The longest reads per genome ranged from 147-260 Kb. An exceptionally long read of 8.26 Mb was obtained from the Zambian line. For each line, RNA was sequenced with short, paired reads (2*150bp) with at least 92M reads/sample, resulting in a total of 8.8-12.3 Mb of RNA sequence per sample (Table 1). *De novo* genomes were assembled with CANU, and the assembly quality was improved following several steps (see Material and Method, supplemental deposit). BUSCO scores, which indicate the completeness of the genome assemblies, ranged from 97.7%(FI) to 99% (ZI) (Table1). The genomes were aligned between lines, and the alignments did not show evidence of major reshuffling, indicating that chromosome sequence identities are mostly consistent across the lines. The total number of singleton genes identified per line ranged from 95.6% (UA) to 98% (ZI) in the final assemblies. The total number of duplicated copies of genes was within the range of 1% (ZI) to 2.1% (TR) . After scaffolding, all genomes contained the 8 chromosomes present in the *D. melanogaster* reference genome (X, 2L, 2R, 3L, 3R, 4, Y, mitochondria). Established genes annotated in the reference genome of *D. melanogaster* were annotated in the 7 new genomes. We detected a total number of genes ranging from 13,471 (ES) to 13,662 (TR), representing 95% to 99% of established genes from *D. melanogaster* (supplemental deposit). The distribution of genes was consistent between lines and chromosomes, while dS values (number of synonymous substitutions per synonymous site) were more variable (Figure 1c, supplemental Table S2.xls), suggesting different selective pressures on genes from each population.

**Table 1:** Descriptive statistics of genome assemblies and annotations

|  | FI<br>(Finland) | DK<br>(Denmark) | ES<br>(Spain) | SE<br>(Sweden) | UA<br>(Ukraine) | TR<br>(Turkey) | ZI<br>(Zambia) |
| --- | --- | --- | --- | --- | --- | --- | --- |
| Estimated DNA base | 9.12mb | 14.49mb | 21.82mb | 15.96mb | 9.66mb | 20.38mb | 10.28mb |
| Estimated genome coverage | 66 | 105 | 158 | 115 | 70 | 148 | 75 |
| Size longest read | 147kb | 220kb | 180kb | 260kb | 160kb | 220kb | 8.26mb |
| Estimated RNA base forward | 8.8mb | 10.7mb | 9.1mb | 9.1mb | 9.2mb | 11.9mb | 9.4mb |
| Estimated RNA base reverse | 9.2mb | 11.3mb | 9.4mb | 9.5mb | 9.4mb | 12.3mb | 9.6mb |
| Complete BUSCO (%) | 97.6 | 98.9 | 98.8 | 98.7 | 97.5 | 98.2 | 98.9 |
| Duplicated BUSCO (%) | 1.2 | 1 | 1.2 | 1 | 1.8 | 2.1 | 0.8 |
| DNA mapping rate (%) | 95.63 | 94.83 | 94.02 | 93.81 | 96.00 | 94.88 | 95.84 |
| RNA mapping rate (%) | 91.83 | 95.87 | 94.36 | 97.11 | 95.29 | 96.11 | 96.05 |
| Genome GC content (%) | 39.16 | 39.31 | 37.98 | 37.61 | 37.70 | 37.86 | 37.83 |
| # annotated genes | 13471 | 13534 | 12594 | 13566 | 13535 | 13662 | 13480 |
| Chromosomes | 8 | 8 | 8 | 8 | 8 | 8 | 8 |

**Figure 1.**
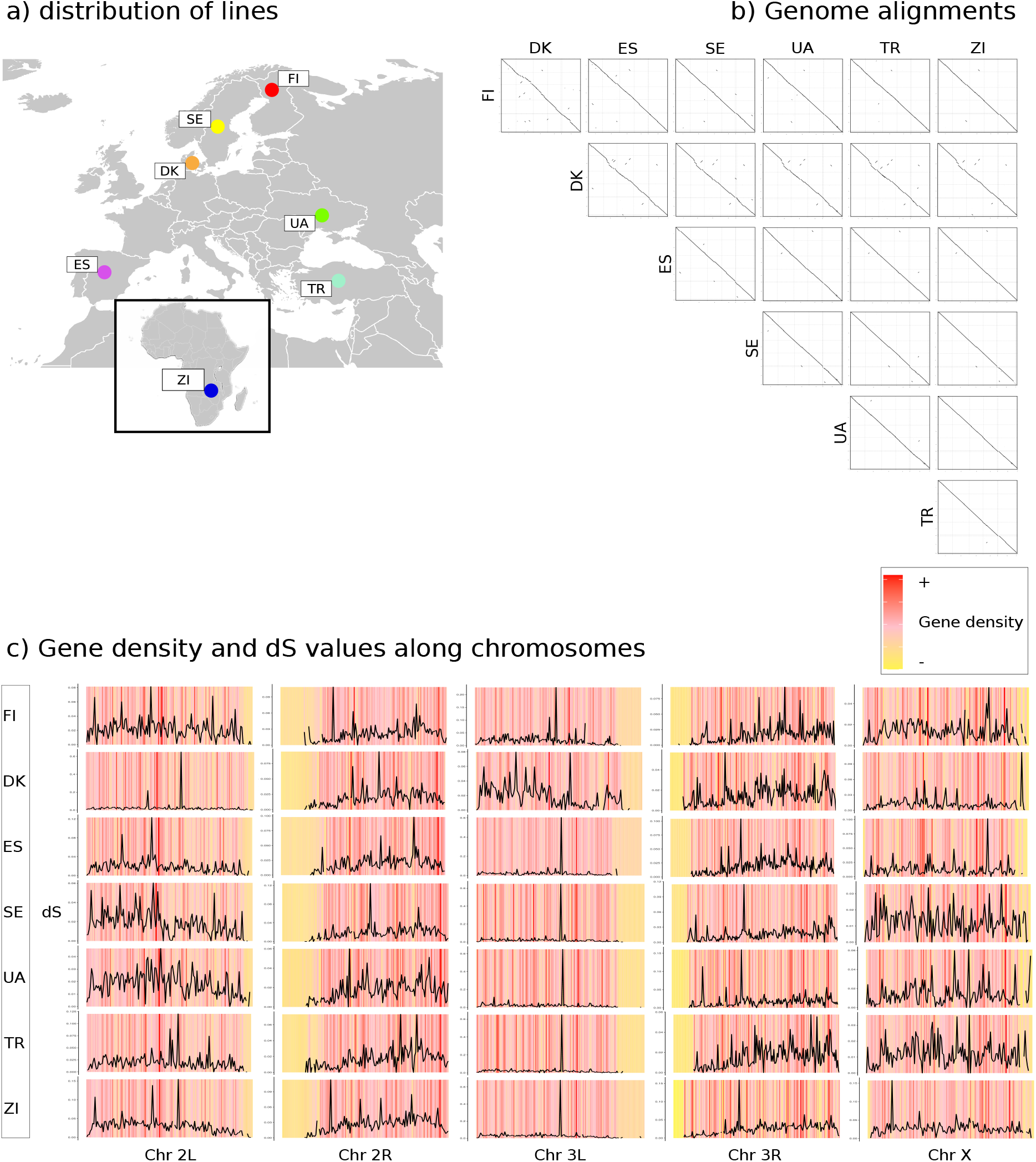
Genome alignments and gene contents. a) Geographic origins of lines; b) Dot plots of all pairwise genome alignments; c) Density and dS values of genes in genomes of each line. The 5 major chromosome arms are shown for each line. The color represent the density of established genes in successive genomic intervals of 160.000 bp. Red denotes highest density, yellow lowest density. Black lines represent average dS values of established genes located in the respective interval. dS was calculated by aligning genes from all lines to orthologs from the reference genomes (see Methods section).

Transcripts were mapped to their corresponding genomes, with percentages of mapping success ranging from 91.23% (FI) to 97.11% (SE) (Table 1, supplemental deposit).

### Proto-genes in D. melanogaster lines

We searched for proto-genes independently in each line of *D. melanogaster*. Proto-genes are here defined as an ORF longer than 90 bp in frame to the direction of transcription, that possesses a putative 5′ and 3′ untranslated region and correspond to a new transcript found in one or more of the 7 genomes, in a non-coding region. These proto-genes show no detectable homology to any species other than *D. melanogaster* nor to established genes of *D. melanogaster* (see methods). On average, we found 1548 proto-genes per line (from 1268 (SE) to 1840 (UA)) (Figure 2a, supplemental Table S3 to S10.xls). The Zambian line did not differ from the European lines in terms of its total number of proto-genes (Figure 2a). Because the Zambian population is known to have a larger effective population size than European populations and to remove deleterious variation more effectively (Kapopoulou et al., 2018), this suggests that the vast majority proto-genes are not deleterious, but instead are effectively neutral with respect to fitness. An average of 593 proto-gene were specific to any one line, with the Zambian line containing more unique proto-genes in proportion (45%) than any of the European lines. This is likely the consequence of the shared out-of-Africa demographic history of the European lines, the relatively high level of gene flow among European populations and the limited amount of European introgression into sub-Saharan African populations (Pool et al., 2012; Kapun et al., 2020). Homology search tools were used to gather proto-genes shared by line in orthogroups. Overall, 5687 orthogroups were established (Figure 2b, supplemental Table S11,S12,S13.xls). Most orthogroups (4010/5687; 71%) contained proto-genes specific to a single line. Among these 4010 orthogroup, the vast majority (3989) comprise a single proto-gene, while the remaining ones comprise proto-genes duplicated within the same genome. The number of orthogroups sharply declines for those containing proto-genes found in two genomes and continues, more moderately, for orthogroups containing instances from 3 to 7 lines. The total number of orthogroups containing a proto-gene shared by all the 7 lines was only 66. As some proto-genes were found to be duplicated and present at different genomic positions within a single line, we investigated the number of copies of a gene contained by an orthogroup (Figure 2c). 16% to 29% of proto-genes were found to be present in at least two copies in the same genome of one line. 35 orthogroups contained over 14 and up to 52 proto-genes, showing the highest level of duplication, even though most orthogroups contained one proto-gene per line. As stated earlier, most of the orthogroups specific to one single line only contained one proto-gene, indicating that most duplications occured in proto-genes that are shared by several lines, and thus are putatively older.

**Figure 2.**
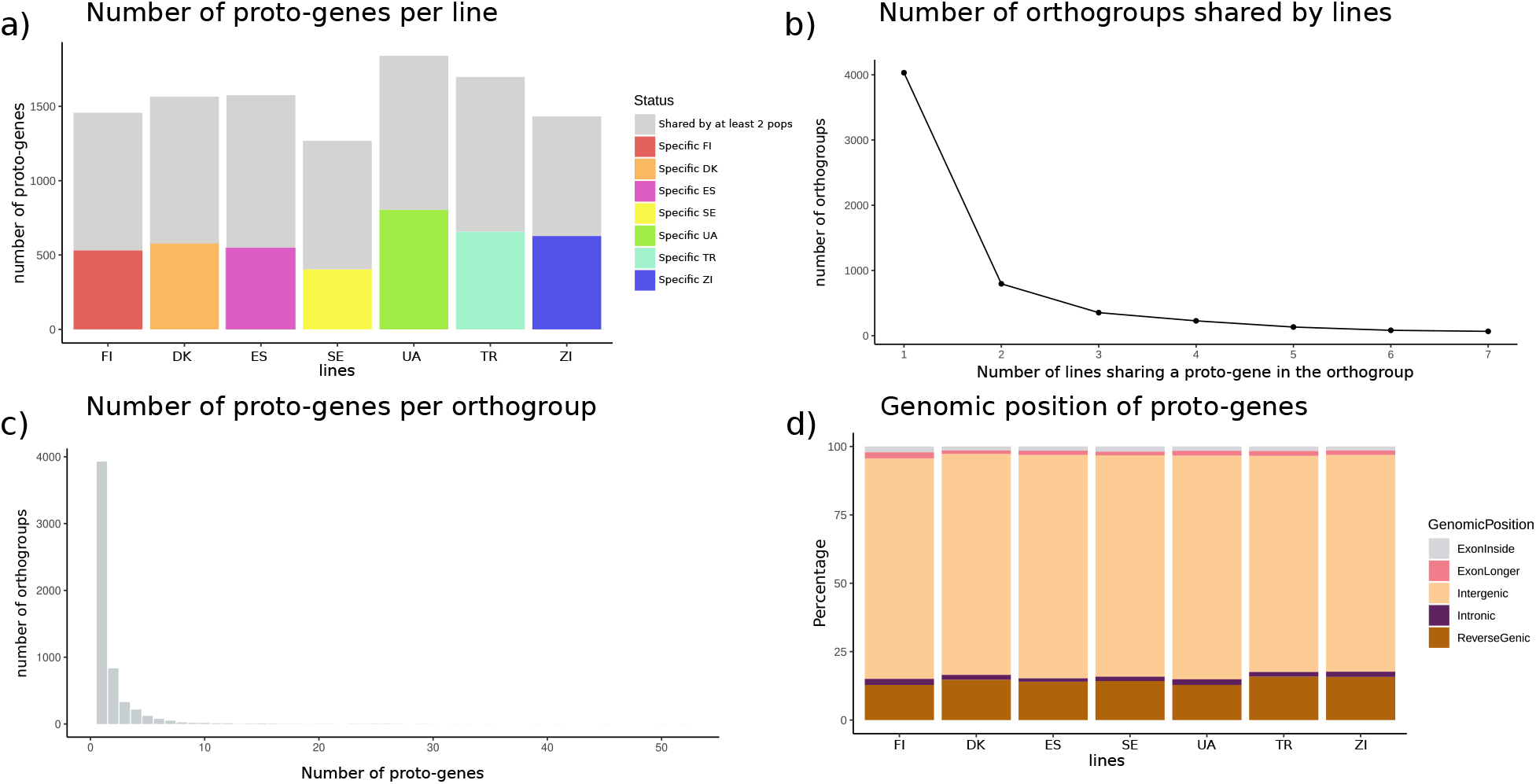
Characteristics of orthogroups from the 7 lines of D. melanogaster. a) Number of proto-genes found in each line. Colors represent proto-genes that are specific to their line. Grey denotes proto-genes with orthologs in other lines. b) Number of orthogroups found in one line or shared by several lines. The X-axis represents the number of lines sharing a proto-gene in an orthogroup. The Y-axis represents the number of orthogroups. c) Number of proto-genes per orthogroup. The X-axis represents the number of proto-genes presents in an orthogroup, the Y-axis represents the number of orthogroups corresponding to these numbers. d) Distribution of genomic positions in which proto-genes are found.

For each line, we next investigated the genomic position of the proto-genes (Figure 2d, supplemental Table S10.xls). The vast majority of proto-genes 79.03% (ZI) to 81.70% (UA), were located in *intergenic* regions. Between 12.80% (UA) and 15.70% (ZI) of proto-genes were found to be *reverse-genic*, i.e. overlapping with an annotated gene but transcribed in the opposite direction. Between 1.47% (DK, ZI) and 2.12% (FI) proto-genes emerged inside an exon of an annotated gene (*ExonInside*). Between 1.21% (DK) and 2.20% (FI) proto-genes partially overlapped to an existing gene and the upstream/downstream non-coding region. Finally, 1.21% (ES) to 2.20% (FI) proto-genes emerged inside an annotated intron. Interestingly, these percentages followed the same trends in all 7 lines.

### Characteristics of proto-genes and their encoded proteins

We next investigated several properties that have been reported to differ between *de novo* genes and established, i.e. evolutionary older genes. First, we compared the length of *de novo* genic ORFs between lines. We find the length of ORFs in proto-genes to be significantly higher for proto-genes which are shared by all 7 lines (average length 235 nt), than for those shared by fewer lines (average 216 nt) with high significance (t-test, p = 1.088e-09) (Figure 3a, supplemental Table S14.xls). Otherwise, no differences in length were found between the proto-genes of the different conservation classes.

**Figure 3.**
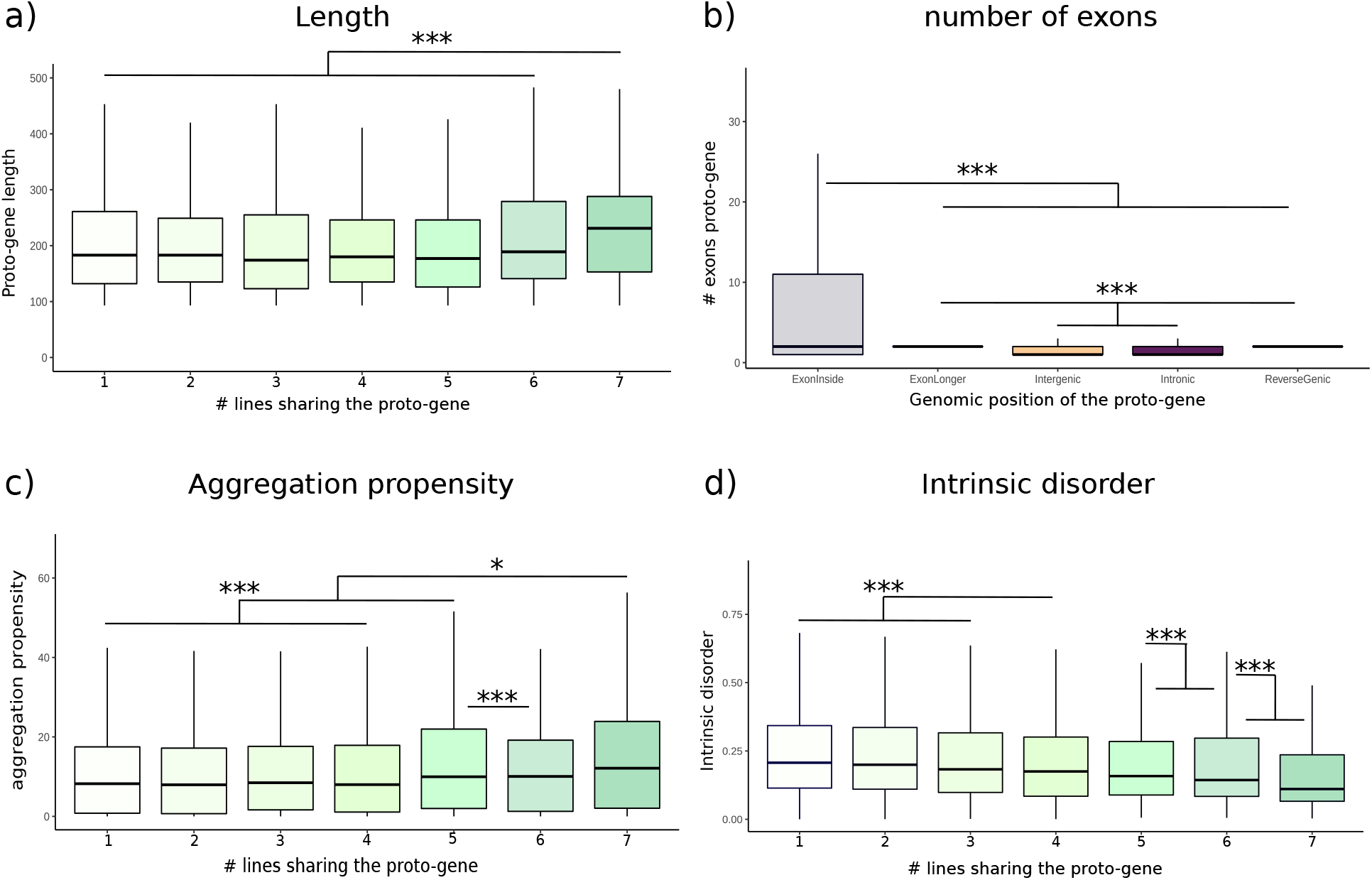
Proto-gene properties. a) Average proto-gene lengths. The X-axis represents the number of lines sharing a proto-gene. The number 1 means that the proto-gene is found only in one line. The Y-axis represents the number of proto-genes. b) Number of exons in proto-genes. The X-axis shows the genomic position of proto-genes. The Y-axis shows the average number of exons. c) Aggregation propensity of proto-genes. The X-axis represents the number of lines sharing a proto-gene. The Y-axis shows the aggregation propensity of proto-genes. d) Intrinsic disorder of proto-genes. The X-axis represents the number of lines sharing a proto-gene. The Y-axis shows the intrinsic disorder of proto-genes.

In each line, around 45% of all proto-genes had introns within their transcripts. (supplemental Table S14,15.xls). We assessed the average number of exons in the unspliced proto-genes, and searched for any correlation between the genomic position of proto-genes and their number of exons (Figure 3b). Interestingly, independently of the lines, proto-genes overlapping an existing exon (“ExonInside”, “ReverseGenic”, “ExonLonger”), had significantly more exons (an therefore introns too) than the proto-genes found in “Intronic” and “Intergenic” positions (supplemental Table S14.xls).

We next investigated if the proto-genes had potential to be translated. Hexamer-scores were all low with average values below 0, as described by (Dowling et al., 2020) (supplemental Table S16.xls). We observed no increase in hexamer score with the number of lines sharing a proto-gene, contrary to earlier reports (Schmitz et al., 2018) that compared genomes across species and, therefore, over much longer evolutionary time scales.

Aggregation propensity and intrinsic disorder of proteins encoded by proto-genes were evaluated and correlated to the number of lines in which the proto-gene was found. The more lines a proto-gene was found in, the more likely its protein was predicted to aggregate (linear model, p = 2e10-16), and the less intrinsic disorder it showed (linear model, p = 2e10-16) (Figure 3c and 3d, supplemental Table S14,17,18.xls). We searched for hydrophobic clusters (HCA) in proteins per proto-gene, since as expected, no annotated domain was detected (supplemental Table S19.xls). We found systematically one HCA cluster in all protein encoded by proto-genes, independent of the number of lines in which they were found (supplemental Table S14,20.xls).

Finally, we compared expression levels between line-specific and wide spread proto-genic ORFs. We find expression to be higher for proto-genes that are found in all 7 lines (49 TPM) than for those shared by fewer lines (average 25 TPM), albeit with much weaker statistical support (t.test, p = 0.0451, supplemental Table S14.xls). As an exception, the expression level of those shared by 5 lines, was higher (139 TPM). Again this is in contrast to earlier studies, e.g. (Schmitz et al., 2020).

### Impact of transposable elements

Transposable elements (TEs) have been repeatedly shown to translocate not only themselves but to carry along other, possibly duplicated genes (Tan et al., 2021) and to integrate in all possible genomic positions, including regulatory regions, exons and introns (Huff et al., 2016; Zhang et al., 2011). TEs are, therefore, widely seen as drivers of evolution. To examine the impact of TEs on the emergence of proto-genes, we tested if proto-genes could be found: (i) inside a TE; (ii) overlapping a TE, or (iii) outside of a TE. We find that between 40% (ES) and 51% (ZI) of proto-genes were found to be outside of a TE (supplemental Table S21,22.xls). 16% to 21% of proto-genes overlapped a TE, suggesting that the insertion of a TE in a non-coding sequence might have provided the sequence changes sufficient to create an ORF. Finally, between 31% and 44% of proto-genes were found to be entirely located inside a TE. These results suggest that the emergence of proto-genes is partly correlated with the presence of a TE.

As 16% to 29% of proto-genes were found to be present in at least two copies in the same genome of one line, we wondered if genes duplicated locally, or rather if the copies were located on different genomic positions. The homologous copies of one single proto-gene were often found on different chromosomes (Figure 4b). Most of the proto-genes found in 2 copies belong to chromosome arms 2R and 3R. Chromosome arms 2L and 3R also share many proto-genes in all lines, except FI. Interestingly, proto-genes found in 2L seem to have homologs in all other chromosomes except the mitochondria, which possesses no proto-genes except in FI. As duplicated copies of proto-genes were mainly found at different genomic locations, we investigated their overlap to TEs. Interestingly, the vast majority of duplicated proto-genes was made up of proto-genes found inside a TE (UA: 74% to ES: 84%) (supplemental Table S22.xls), and a small amount overlapped to a TE (DK: 6% to UA: 11%). This result strongly suggest that transposable elements were responsible for most duplication events in duplicated proto-genes, and again demonstrates the importance of TEs in the origin of proto-genes, and their spread within the genomes. We next calculated the frequencies or TEs and of proto-genes per 160,000 kb intervals on all chromosomes (figure 4c). TEs were detected in each of the 7 assemblies, with proportions ranging from 15.64 to 19.31% of a genome′s content (supplemental Table S23.xls). The 7 genomes contained LINEs, low complexity families, LTR elements, satellites and simple repeats, SINEs. The TE distribution was highly biased along chromosomes with the highest number of TEs detected at telomeres. Their distribution was the opposite that of established genes (Figure 1a), which were mainly absent in telomeres. Somewhat unexpectedly, proto-genes were distributed regularly along chromosomes, and were also present in telomeres. Therefore, the distribution of proto-genes is less biased than the distribution of TEs: high density regions of TEs did not correspond to high density regions of proto-genes.

**Figure 4.**
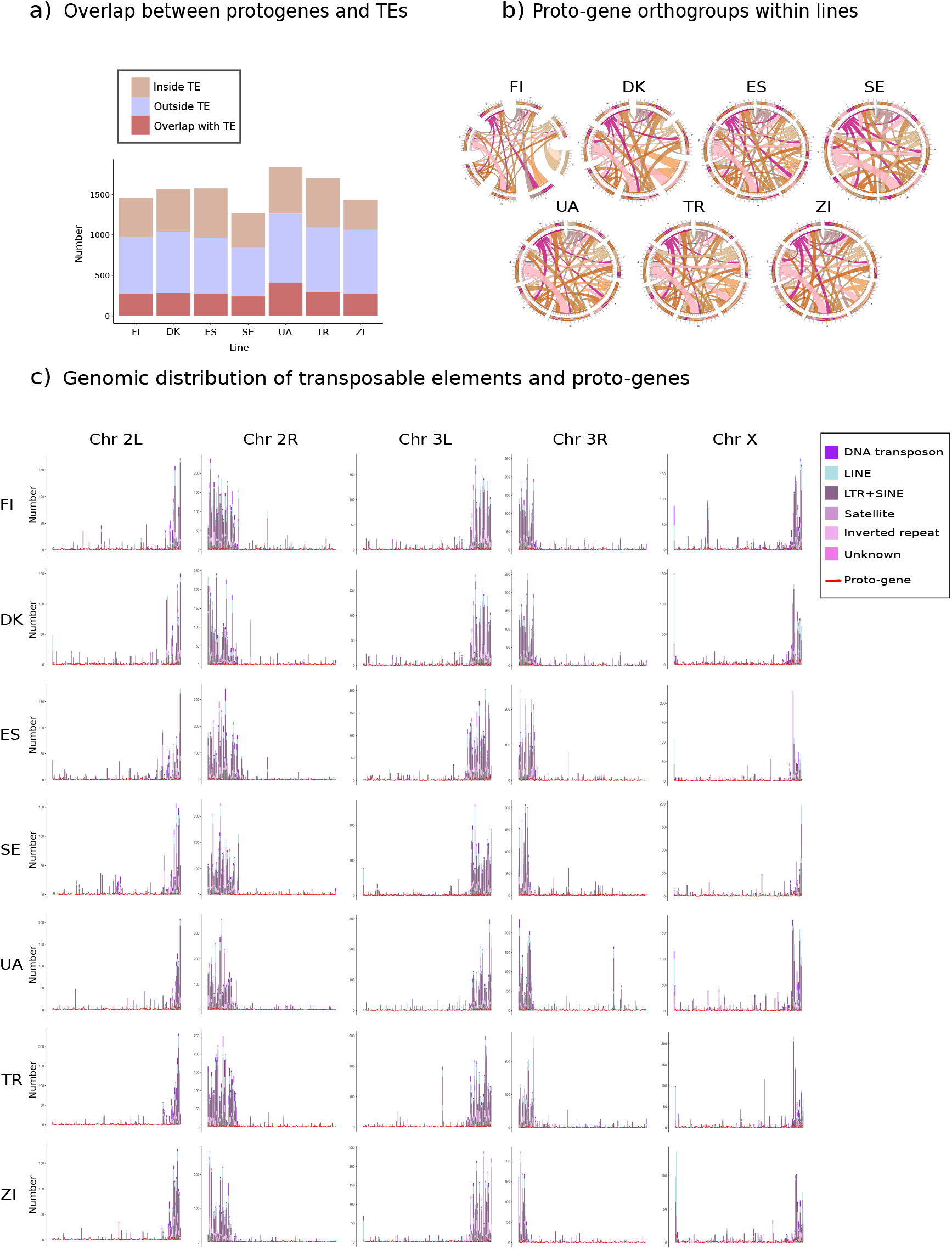
Proto-genes and transposable elements. a) Overlap between proto-genes and transposable elements inside lines. The x axis represents the lines and the y axis represents the number of proto-genes. A brown color represents proto-genes that are located inside of a transposable element; the violet color represents proto-genes that do not overlap with a transposable elements, the pink color represents proto-genes overlapping partially with a transposable element, but not entirely. b) Circle plots representing the genomic location of proto-genes duplicated inside a line’s genome. A circle’s borders represent chromosomes. c) Distribution of TE (bars) and proto-genes (red lines) along the 5 major chromosome arms. The X-axis represents chromosomes, in successive intervals of 160.000 bp. The Y-axis represents the total number of elements (TE or protogenes) present in each interval

### Non-coding homologs of proto-genes

We next attempted to identify, for every proto-gene in every lineage, homologous non-coding sequences in those lines in which no coding copy (i.e. homolog) of the respective proto-gene (termed “query” hereafter) had been found. For this, we used a pipeline (see Methods, github link) involving BLAST and synteny detection. This would enable us to locate the homologous sequence derived from the query proto-gene and identify sequence differences (“enabling mutations”, see also (Zhang et al., 2019)) which enabled the emergence of the proto-gene in first place.

First, proto-genes shared by all lines were removed from the analysis. A total of 5,613 query proto-genes (out of 5,687 orthogroups, see above) of which 3,499 were without introns and 2,114 were with introns, were investigated. Each proto-gene was used as a query to detect non-coding homolog in all other lines in which the proto-gene was missing, (searched in an average of 5.5 lines), giving rise to a total of 30,606 homology searches. Restrictive homology searches were performed initially against syntenic regions of the target lines, and then in full genomes when no hit was found. When several hits for a query were found in a single line, only the best BLAST hit was kept for further analysis and identified as the “non-coding homolog”.

94.9% (DK-FI) to 99.8% (ES-ZI) of all proto-gene queries were found to have non-coding homologs in one or several target lines (Figure 5a), resulting in 27,699 non-coding homologs (90.5% of the search). Sequence identities between query proto-genes and their target non-coding homolog were between 80% to 100%, indicating a high reliability in the identification of non-coding homologs. 43.1% of the alignments between queries and their non-coding homologs had a sequence identity ranging from 95% to 100%, based on the Levenshtein ratio (Figure 5b).

**Figure 5.**
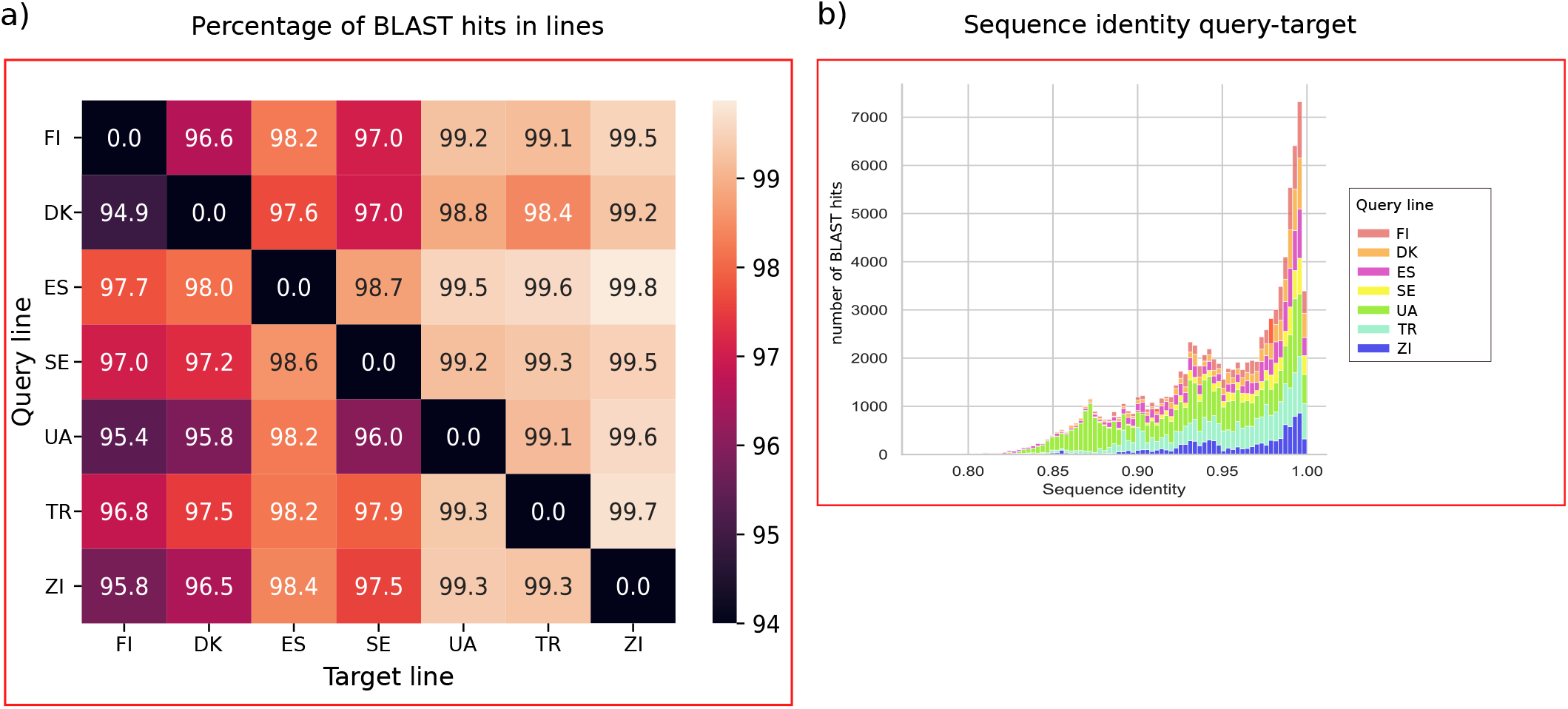
Detection of non-coding homologs by homology searches. a) Percentage of BLAST hits between query proto-genes of each line and target lines. b) Percentage of sequence similarity between proto-genes and their “non-coding homologs”, according to the Leveinshtein ratio. Each color represents a line.

### Enabling mutations

The 27,699 non-coding homologs were aligned to their homologous proto-genes, after computationally splicing the proto-genes that contained an intron. The alignments were analysed to determine the enabling mutations that distinguish proto-genes from their non-coding homologs. Among the 27,699 non-coding homologs, 23,159 had at least one mutation that can be identified as rendering it non-coding. The remaining 4,540 had no mutations. Their existence might be explained by the presence of a longer ORF overlapping with them, hiding a smaller ORF (supplemental Table S24.xls). These non-coding homologs were removed from further analyses.

We determined six different features which might explain the different coding characters between queries and non-coding homologs: i): absence of a start codon; ii): absence of a stop codon; iii): insertion or deletion causing a frameshift mutation; iv): presence of a premature stop codon or downstream start that would cause the encoded protein shorter; v): different sequence coverage in the alignment, causing the end or the beginning of the non-coding homolog different from the proto-gene. This may results from, for example, the insertion of a TE in either of the two sequences; and vi): Absence of a detectable transcription event.

Among these six features, the lack of transcription was the most common difference between a noncoding sequence and its homologous proto-gene (Figure 6). Of the non-coding homologs, 85% were not transcribed. Furthermore, 39.5% of the non-coding homologs have a frameshift mutation compared to their proto-genes, which would result in a major change in the protein sequence and size, if any protein is encoded. A premature stop codon was present in 25.35% of the non-coding homologs, resulting in a shorter ORF than their proto-gene. Finally, 9.18% of the non-coding homologs have no stop codon, 9.03% have no ATG start codon, and 8.83% display a beginning or end of sequence which does not correspond to the proto-gene (Figure 6). Most often, several of these feature co-occurred in a non-coding homolog, explaining why the sum of all percentages exceeds 100.

**Figure 6.**
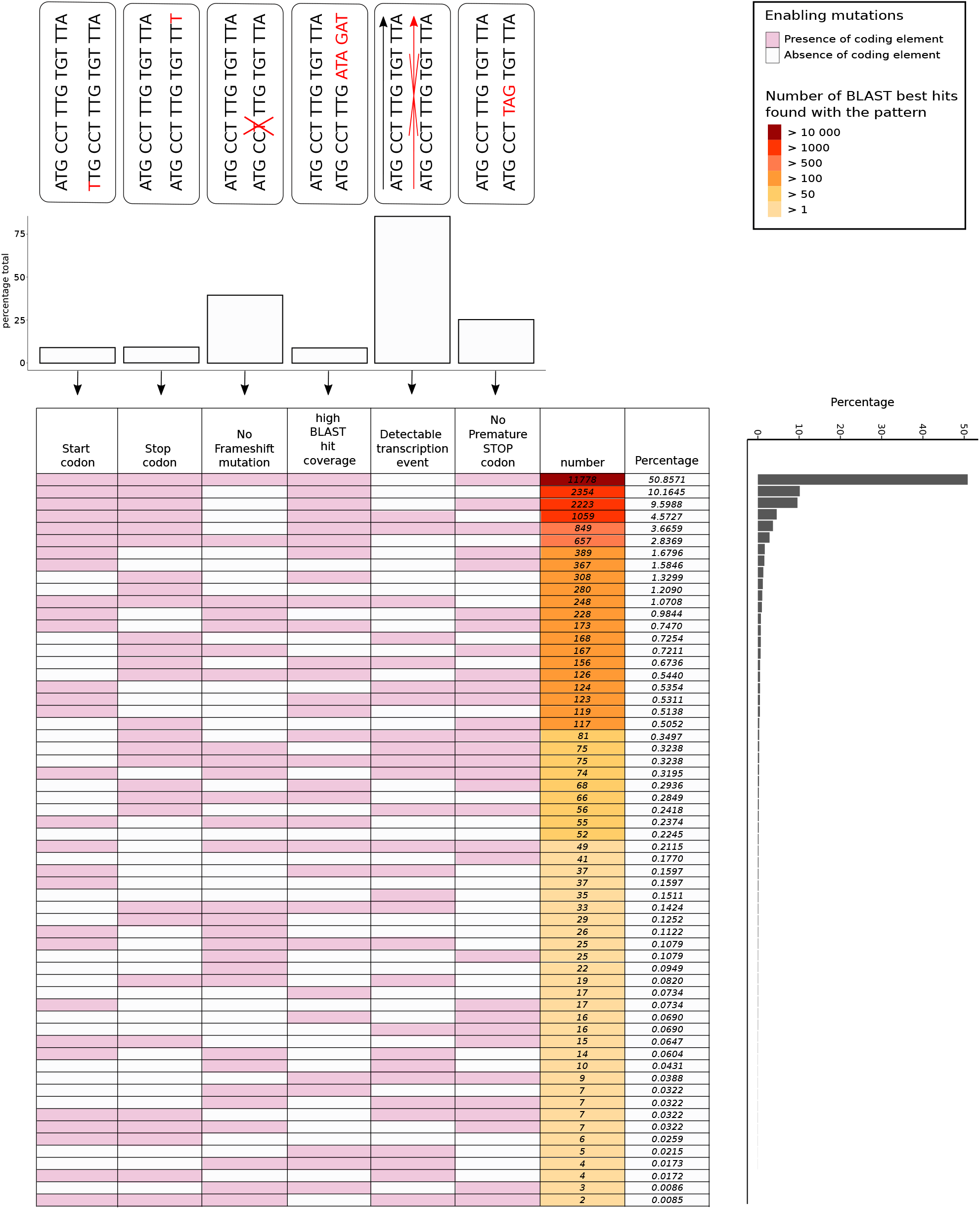
Enabling mutations between proto-genes and their “non-coding homologs”. The barplot on the top of the figure represents the percentage (y axis) of 6 elements (x axis) that are not presents in “non-coding homolog”. The table represents the percentage of each combination of enabling elements. The pink colors indicates that the considered element (ex Start codon), is present in the non-coding homolog. The grey color indicates that the elemetis not present in the non-coding homolog.

We next studied the combination of enabling features observed in non-coding homologs. 60 combinations of these 6 events were observed, some being represented by one single event. The most frequently observed combination observed in proto-genes was one single event: a single lack of transcription event, which was the case for 11,778 non-coding homologs. A frameshift mutation combined with a lack of transcription event was the second most common combination, and was found in 2,223 non-coding homologs. Furthermore, frameshift mutations combined with a premature stop codon, and frameshift mutations combined with the absence of transcription plus presence of a premature stop codon often occurred together (1,059 and 2,354 times, respectively). The absence of start and stop codons were the least frequently observed events. Overall, when comparing the presence of an ORF and a detectable transcription event, 50.86% of all non-coding homologs have all of the coding elements in the ORF but are not transcribed, whereas 14.53% of all non-coding homologs are transcribed but do not possess all the coding elements of an ORF. Finally, 34.61% of non-coding homologs do not have a coding ORF and are also not transcribed.

We investigated in further detail the nature of the mutations that constitute the difference between protogenes and their non-coding homologs (supplemental Table S24,25.xls). 36% of non-coding homologs that have no start codon have 0 ATG elements. The presence of two substitutions in the ATG was widely observed (52%). One substitution in the ATG was observed in 20% of all cases. We observed the same trend with the stop codons, in which all three nucleotides constituting the homologous stop codon were substituted in (25%) of all cases.

When a frameshift mutation was observed, we studied how many nucleotides were missing or added in the non-coding homolog. The most frequent frameshift was due to only one insertion or deletion. Finally, we studied in further detail the absence of a transcription event in non-coding homologs. The absence of a transcription event could be attributed to three different situations: i) no transcript was found; ii) a transcript was found but in the reverse direction to the ORF; or iii) a transcript was present (in the forward or reverse direction) but did not cover the entire ORF. 50% of the non-coding homologs were located in a transcribed sequence, but the transcript was in opposite direction to the predicted ORF, even in cases when all coding elements were present. For 10% of the non-coding homologs, a transcription event was observed, but did not cover the length of the entire transcript, in either the forward or reverse direction. Furthermore, 40% of the non-coding homologs were not transcribed at all.

## Discussion

In the present study, inbred lines from 7 populations of *D. melanogaster* were sequenced and their genomes and transcriptomes were assembled. We detected proto-genes and studied their properties and mechanisms of emergence. Our methodology allowed us to detect very young proto-genes that emerged within lines of a single species using a common sequencing and annotation framework. This study thus fills the gap between earlier phylogenetic studies that used long evolutionary distances, and population level studies that either used sparsely annotated reference genomes or did not investigate the genetic mechanisms underlying the creation of proto-genes. A major result of our work is that proto-gene emergence is a frequent event, even when viewed from the perspective of populations.

### Proto-gene birth and death

Interestingly, the average number of proto-genes found per line (around 1,500 proto-genes) is in a similar range of numbers to that reported in other species. For example, in yeast, 1,900 candidate proto-genes were detected in *S. cerevisiae* for a much smaller genome and for much longer evolutionary divergence between species (Carvunis et al., 2012). In humans, Dowling et al. (2020) identified 4429 *de novo* transcribed ORFs specific to humans. Since most orthogroups (71%) detected in our study contained proto-genes specific to a single line, we conclude that there is a rapid turn-over within the species. Such a pervasive creation and rapid turn-over of proto-genes has already been suggested by cross-species comparisons (Schmitz et al., 2018; Prabh and Rödelsperger, 2022) but not yet demonstrated on evolutionarily short time scales. Furthermore, the fairly equal distribution of proto-gene′ numbers across lines suggests a neutral birth-death process, at least over such short time scales. The comparability of our numbers with results from cross-species studies, also including those in *Drosophila* (Heames et al., 2020), indicates that such a neutral regime extends to time scales spanning at least several speciation events.

62 proto-genes were present in all 7 lines and might be considered as established *de novo* genes. Again, the respective counts are comparable to earlier studies using different approaches: Heames et al. (2020) found 41 putative *de novo* genes that emerged specifically in *D. melanogaster*, while Zhou et al. (2008) reported 72 *de novo* genes and Zhao et al. (2014) identified 142 segregating and 106 fixed testis-expressed *de novo* genes in *D. melanogaster*.

### Proto-gene properties

In each line, around 80% of the detected proto-genes emerged in intergenic regions, while the next largest group (15%) emerged on reverse strands of other genes. These results are in line with earlier cross-species studies of humans (Grandchamp et al., 2022; Dowling et al., 2020), which showed that older *de novo* genes are more often exonic, while younger ones are more likely to be found in intergenic regions. Again, this result strongly suggests a neutral process of proto-gene emergence and loss in the early stages of their formation. It seems intuitive that proto-genes can emerge more easily in non-coding regions, as established genes are constrained by purifying selection (Finseth et al., 2014), whereas non-coding regions are less constrained by selection (Knibbe et al., 2007). Moreover, a proto-gene overlapping with an existing gene would probably have a lower likelihood of gaining an adaptive function compared to one emerging in a non-coding region.

When comparing properties of lineage-restricted proto-genes to those that are wide spread, we found several notable differences. Compared to established genes, proto-genes were shorter (235nt vs. 218nt) as is known from many cross-species comparisons. Somewhat surprisingly, we find that, even at such a short evolutionary time scale, fixed proto-genes (those present in all 7 lines) were longer than less pervasive ones. In each line, almost half of all proto-genes contained introns, with the average number of introns being highest in exonic proto-genes, and lower in intergenic and intronic proto-genes. This finding strengthens recent reports showing that *de novo* genes can contain introns (Wu et al., 2011; McLysaght and Guerzoni, 2015; Zhang et al., 2019; Grandchamp et al., 2022), and demonstrates that introns can appear at the earliest stages of proto-gene emergence.

We observed that the computationally predicted amount of intrinsic disorder in proteins encoded by proto-genes decreased with their pervasiveness, and thus with their relative age, while their aggregation propensity increased. Since this prediction aligns with some studies (Dowling et al., 2020) but disagrees with others (Schmitz et al., 2018), this trend deserves further attention in experimental follow-up studies. Furthermore, we found no increase in formation of hydrophobic clusters with age. The latter process would support the idea that proto-genes encoding novel proteins become more structured and compact over time but do not play a role in the early stages of proto-gene emergence.

### Enabling mutations

Our approach allowed us to track enabling mutations and the features associated with proto-gene birth by mapping them precisely to homologous regions in other genomes. Our homology-based searches retrieved the majority (90.5%) of all non-coding homologs with high levels of sequence coverage and identity. Contrary to most earlier studies, these assignments are non-ambiguous with near perfect sequence identities. Even for most of the proto-genes that have been translocated within chromosomes or to other chromosomes, non-coding homologs could be identified, due to the short divergence times under consideration. While earlier studies have reached discordant conclusions regarding the temporal order of gain of transcription vs. gain of ORFs (Schlötterer, 2015), we here were able to identify and quantify the frequency of features (or combinations thereof) that distinguish proto-genes and their non-coding homologs. In our study, we discovered that transcription is the feature most frequently missing in non-coding homologs. About half or all non-coding homologs have all ORF specific features but lack detectable and/or complete transcription. Of the remaining half of non-coding homologs, only 14.53% are fully transcribed, but lack all features of an ORF. About one third have neither a complete ORF, nor are they transcribed. Taken together, these observations suggest that transcription status change very rapidly. Therefore, the presence of an ORF is not a sine-qua-non prerequisite for proto-gene emergence, but more often by chance may precede the gain of transcription than the other way around.

The rapid turnover of pervasive transcription has been widely documented in recent years (Neme and Tautz, 2016; Wade and Grainger, 2014; Clark et al., 2011). Underlying mechanisms (reviewed e.g. in (Kim and Jinks-Robertson, 2012)) include epigenetic events such as methylation turnover (Ginno et al., 2020), replication and spatial chromatin changes (Candelli et al., 2018), the birth and death of promoters or their interconversion into enhancers (Majic and Payne, 2020) and their putative bidirectional functioning (Young et al., 2015). Antisense transcripts occur frequently (Pelechano and Steinmetz, 2013) by exploiting transcription loops of genes transcripts. Indeed, among the 17% proto-genes overlapping genic sequences in our data set, 15% were reverse-genic to established genes, suggesting that these proto-genes emerged as antisense transcripts. These data also lay the foundation for further in-depth investigations into the mechanisms of new transcripts formation and the causes of their transient behaviour. Furthermore, the here reported abundance of mainly short ORFs in non-coding DNA makes it likely that many are randomly translated (Ingolia et al., 2014). Indeed, the coding potential of thousands of small ORFs in Drosophila has been demonstrated using ribosomal profiling (Aspden et al., 2014; Patraquim et al., 2022), and pervasive transcription has also shown to evolve new functional protein (Ruiz-Orera et al., 2018). Overall, we observed all possible combinations of features to be involved in the creation of new ORFs, implying that there are many different paths giving rise to new ORFs. Interestingly, 10% of non-coding homologs lacked all six features, showing that evolution can quickly build ORFs during population divergence.

Finally, we considered the criterion “different size” as one of the 5 criteria to define the appearance of an ORF. It applies when the end or the beginning of the sequence of the proto-gene and its non-coding homolog diverge completely. This can be due to genome reshuffling between lines, or an error in the genome assembly. It also can be explained by the insertion of a TE.

### Transposable elements

Growing evidence has shown that TE insertions enabled the emergence of protein-coding genes or non-coding RNAs, and that some of them acquired essential functions over the course of evolution (Joly-Lopez and Bureau, 2018; Naville et al., 2016). Among the most famous examples of new genes that emerged thanks to transposable elements, are rag1 and rag2, which are involved in the vertebrate immune system (Huang et al., 2016; Kapitonov and Koonin, 2015), and the syncytin genes of mammals, which contributed to the emergence of the placenta (Malik, 2012). When looking at the features enabling protogene emergence from non-coding homologs, only 9% of non-coding homolog showed strong reshuffling of the end or the beginning of their sequence. This suggests that only 9% of the non-coding homologs gained an ORF via the insertion of a TE in their sequence. TEs are thus important, but not the dominant driving force in the creation of novel proto-genes.

However, once emerged, a high number of proto-genes were found to be associated with a TE, either by being located inside a TE (25-30%), or by overlapping with a TE (15-20%). Indeed, even though TE insertions are not the main events enabling proto-gene emergence, their presence in genomes may promote other mutations that contribute to proto-genes emergence. TEs are know to be enriched in genomic hotspots, which are often concentrated in telomeres (Venner et al., 2009; Robillard et al., 2016), and this fact was confirmed by our results. TEs that insert in genic regions are much more likely to be shaped by selection (Catlin and Josephs, 2022), and may be quickly eliminated from genomics regions that are under strong purifying selection (Bourque et al., 2018). We found that proto-genes are distributed regularly across chromosomes. Consequently, proto-genes overlapping with TEs are more likely to overlap those more subject to selection than to telomeric TEs, which might explain they high potential of mutation. Activated transcription was found to be the most frequent features associated to proto-gene emergence, and can also be due to the proximity of proto-genes to TEs, as TEs have been shown to contribute and modify cis-regulatory DNA elements and modify transcriptional networks (Bourque et al., 2018; Jacques et al., 2013; Trizzino et al., 2017; Bejerano et al., 2006; Bourque et al., 2008; Lippman et al., 2004; Wang et al., 2015).

Finally, among the average 20-25% of proto-genes that were duplicated inside a line, around 80% were located inside of a TE. Thus, TEs appear to play an important role in duplicating proto-genes, and are able to translocate them within genomes. The ability of TEs to effect rapid translocation has already been reported at the population level (Kofler et al., 2015; Bartolome and Maside, 2004), but to the best of our knowledge, this is the first time that they have been reported as a duplication vector of proto-genes.

## Methods

### Genetic material

To get a representation of the genetic diversity in Europe, we sequenced genomes of isofemale lines collected from 6 different European populations (supplemental Table S1.xls): Finland (FI), Sweden (SE), Denmark (DK), Spain (ES), Ukraine (UA) and Turkey (TR), and one isofemale line from the presumed ancestral range in sub-Saharan Africa Zambia (ZI), which diverged from the European populations around 14000 years ago (Laurent et al., 2011; Li and Stephan, 2006) The lines were established as isofemale lines with 5 generations of single sib-sib inbreeding, then subsequently maintained in vials with *en masse* sib-sib mating of 50-100 individuals per generation.

### Nucleic acid extraction and sequencing

High molecular weight genomic DNA was extracted from pool of 100 individuals (mixed female and male) from each line with the DNA Easy Blood and Tissue Kit (Quiagen, Hilden, Germany). RNA contamination was removed using chloroform extraction and isopropanol precipitation, and quality control was performed using agarose gels, a UV spectrometer, and *NanoDrop* measurements. For each line, DNA libraries were prepared with the Ligation Sequencing Kit LSK109 and the Flow Cell Priming Kit EXP-FLP001. Long-read DNA sequences were generated using Nanopore sequencing, with flow-cells 106D. The reads were generated in FAST5 format, and stored for each line. The line from Zambia was sequenced twice, since the first round of sequencing did not generate enough reads.

Since our goal was to capture as many novel transcripts as possible, irrespective of their temporal, developmental or spatial expression profiles, we extracted total RNA from a pool of 5 individuals (2 adult males, 2 adult females and 1 larvae) from each iso-female line. To minimise the number of missed *de novo* transcripts, we aimed at high coverage. Note that missing some weakly expressed transcripts will only slightly influence our results: we are interested in the comparison between stably expressed *de novo* ORFs, their presumed ancestors and their relation to random sequences and just any ORF with transcription *potential* (i.e., whether transcribed or not). Total RNA following protocols widely used for *D. melanogaster* (Trizol based) (Emery, 2007). DNA contamination was removed with DNase treatment, and control quality (agarose gels, UV spectrometer, NanoDrop measurements) was performed. The RNA samples were sent to the Institute of Clinical Molecular Biology of Kiel for sequencing. Libraries were constructed for Truseq stranded RNA, and RNA was sequenced with RNA SeqNova6000 S1 2*150bp, with 92M reads/sample.

### Sequence preparation and de novo genome assembly

For each line, the long DNA reads generated were converted into fastq format with guppy basecalling from the Oxford Nanopore Technologies’ basecalling algorithms (https://nanoporetech.com/). Identified doublons were removed from the datasets. Reads shorter than 50bp were removed from the assembly with Filtlong (github rrwick/Filtlong). RNA reads were trimmed with Trimmomatic to remove the adaptors AGATCGGAAGAGCACACGTCTGAACTCCAGTCA in forward and AGATCGGAAGAGCGTCGTGTAGGGAAAGAGTGT (Bolger et al., 2014). The quality of reads was assessed with FASTQC (Wingett and Andrews, 2018).

For each line, DNA reads were converted into FASTA format with seqtq (Anaconda package), and assembled with CANU (Koren et al., 2017) with 137Mb as a reference genome size from BDGP6.28 genome in Ensembl (Yates et al., 2020). The quality of the assembly was assessed with BUSCO with insecta odb9 as a reference database and *fly* as reference species (Manni et al., 2021).

To improve the quality of the assembly, the contigs were first polished with the RNA reads specific to each line. The RNA reads of each line were mapped on the contigs of their respective assembly (generated previously with CANU), with the spliced aware software Star (Dobin et al., 2013). The sam files were converted to BAM, sorted and indexed with samtools programs (Danecek et al., 2021; Bonfield et al., 2021). The resulting genomes were polished with Pilon (Walker et al., 2014). The quality of the new genome assemblies was assessed with BUSCO.

Subsequently, the quality of the assemblies was further improved by mapping the long DNA reads of each line to their respective reference genome with Minimap2 (Li, 2018). The resulting sam files were converted to BAM, sorted and indexed with samtools. As the 7 lines were inbred for at least 5 generations, the assembled genomes had low levels of heterozygosity. Nevertheless, we resolved haplotypes with purge-haplotype programs using the readhist, contigcov and purge options (Roach et al., 2018). The resulting files were polished with Pilon (github broadinstitute/pilon). The quality of the new contigs assembly was assessed with BUSCO. Finally, the contigs were scaffolded with the reference genome BDGP 6.28 of *D. melanogaster*. Scaffolding was performed with RaGOO/Ragstat (Alonge et al., 2019). The quality of the assembly was assessed with BUSCO.

### Genome annotation and transcriptome assembly

In the 7 genomes, repeats were modeled and classified with RepeatModeler (Flynn et al., 2020), and masked with RepeatMasker (http://www.repeatmasker.org). Homology-based methods were used to annotate the genomes with the software GeMoMa (Keilwagen et al., 2019). At this stage, we only aimed to annotate the genes already known and annotated in the reference genome of *D. melanogaster*. Reference genes were used as a support for gene detection and exon/intron structure detection. The BDG releaseP 6.28 of *D. melanogaster* was used as the reference genome with matching annotation data, and tBLASTn (Altschul et al., 1990) was used for homology searches. Pairwise alignments of genomes were performed between lines with Chromeister (P′erez-Wohlfeil et al., 2019). All established genes were retrieved in lines, aligned to the reference annotated genes with mafft (Katoh et al., 2002) and pal2nal (Suyama et al., 2006), and dS values were calculated with python program using Tajima′s formula. dS values correspond to the number of synonymous substitutions per synonymous site, in our case between each gene from lines and their homologous reference gene.

For each line, respective RNA reads were mapped on their reference genome with Hisat2 (Zhang et al., 2021), using the splicing aware option. The resulting SAM files were converted to BAM format with Samtools, and the BAM’ files were sorted and indexed. For each line, transcriptomes were assembled with Stringtie (Pertea et al., 2015). FASTA, GTF and GFF files for transcriptome assemblies were retrieved with TransDecoder (https://github.com/TransDecoder).

### Detection/assessment of proto-gene status

In each assembled transcriptome, the TPM (Transcripts Per Million) value was calculated for each transcript. Following earlier studies and statistical evaluations of transcript persistence (Schmitz et al., 2018; Dowling et al., 2020), transcripts with TPM value inferior to 0.5 were removed. ORFs in the remaining transcripts (*trORFs*) were detected with the software GetORF from EMBOSS (Rice et al., 2011) with a size ranging from 30-3000 amino acids, in forward direction of the transcript. Transcript names and the ORF’ positions in the spliced transcript were assessed. `GetORF` considers the end of a transcript as a stop codon, and indeed some false-positive ORFs were detected by the software. These ORFs were removed from the dataset, as only ORFs starting with a start codon and ending with a stop codon were taken into account.

In order to remove known annotated proteins from the ORF dataset, ORFs were translated into proteins, and used as a query for a protein BLAST search against Drosophila proteomes in the forward direction of transcription. Proteomes of the 11 *Drosophila* species available in the Ensembl database were downloaded as BLAST targets: *Drosophila ananassae, Drosophila erecta, Drosophila grimshawi, Drosophila mojavensis, Drosophila persimilis, Drosophila pseudoobscura, Drosophila sechellia, Drosophila simulans, Drosophila virilis, Drosophila willistoni* and *Drosophila yakuba*. The reference genome of *D. melanogaster* was also included. All tORFs with no hit at an e-value of 10-2 were stored as putative *de novo* tORFs. To refine the search, a second BLAST was performed against the proteomes of 15 dipteran species present in Ensembl and covering all dipteran subfamilies: *Anopheles albimanus, Anopheles culicifacies, Anopheles darlingi, Anopheles farauti, Anopheles gambiae, Glossina austeni, Glossina morsitans, Lucilia cuprina, Lutzomyia longipalpis, Mayetiola destructor, Megaselia scalaris, Musca domestica, Phlebotomus papatasi, Stomoxys calcitrans* and *Teleopsis dalmanni*.

All transcripts whose tORF had no BLAST hit were considered as proto-genes. For transcripts containing several ORFs with no BLAST hit, only the longest tORF was retrieved and stored as “putative” proto-gene. The genomic positions of the proto-genes were determined with custom python script. Genomic positions of unspliced and spliced proto-genes were retrieved by combining the GTF file information and the position in the spliced transcript of the proto-genes’ ORFs. Each position was verified by reconstructing the ORF in the FASTA genomes. For each putative proto-gene, the transcript’s name, number of exons present, genomics position (of the unspliced transcript), direction of the transcription (forward or reverse), TPM, FPKM and coverage values were extracted and normalised. The transcripts that overlapped and might represent the alternative splicing of one single gene were also annotated.

Bedtools (Quinlan and Hall, 2010) was used to attribute a “Position Category” to each proto-gene. Five “Position Categories” were established: Overlapping with an intergenic region, overlapping with an intron, inside a gene in frame, overlapping with a gene in reverse direction, and overlapping with a gene and the surrounding intergenic region.

### Building orthogroups of proto-genes between lines

Proto-genes were identified in each of the 7 lines of *D. melanogaster*. To identify proto-genes that were common to several lines, and proto-genes that were paralogs in one single line, proto-genes from all lines were pooled together and scanned for orthogroups with the software `Orthofinder` (Emms and Kelly, 2019) (supplemental Table S26.xls).

### Characteristics of proto-genes

Size, GC content and the level of RNA expression (TPM and FPKM) were assessed for each proto-gene. For translated protein sequences, intrinsic disorder was assessed with Iupred (http://iupred.enzim.hu). Aggregation propensity was assessed with TANGO (Fernandez-Escamilla et al., 2004). The presence of annotated domains was searched with Pfam scan from PFAM database (Mistry et al., 2021), and the presence of new hydrophobic clusters was searched for with SEG-HCA (Faure and Callebaut, 2013). The overlap of proto-genes with transposable elements was investigated with custom scripts. Some orthogroups contained several proto-genes from the same line, originating from different regions of the genome. We investigated the duplication events within genomes, and mapped the TEs and proto-genes distribution for each chromosome.

### Assessing mechanisms of proto-gene birth

Proto-genes absent from at least one line were used as the query to search for homologous non-coding sequences in the lines in which they were not detected, in order to study putative enabling mutations to a coding state. A python pipeline was designed to find non-coding homologs and investigate mutations further (supplemental Table S27.xls).

In each query orthogroup, the unspliced sequence of the proto-gene was retrieved. For orthogroups that contained several orthologs, when the orthologs were not 100% identical, the sequence which showed the highest similarity to the other ones was selected. When the proto-gene contained an intron, the intronic sequence was lowered in the FASTA file, and the exonic sequences were left in upper letters (supplemental Table S28,29.xls). Proto-genes containing an intron and whose unspliced size was greater than 10,000 bp were removed from the analyses, as such long sequences would have biased the homology search. “Query proto-gene” hereafter refers to proto-gene under investigation. “Target line” refers to the line in which a non-coding homolog is searched for.

### Step 1. Identifying syntenic regions in target lines

The two established genes neighbouring the query proto-gene were identified in the target line. Corresponding genes were looked for in the GTF file of the target line, and the DNA region between these two genes was stored as “Target syntenic region”. Whenever one of the two genes was missing, the second next surrounding gene was used. When a proto-gene was not surrounded by two genes but only by one gene and the end of the chromosome, the corresponding region was retrieved in the target line. When the two target surrounding genes were not found in syntenic region in the target line, the whole genome was used as a target.

### Step 2. Detecting non-coding homologs

The unspliced sequence of query proto-gene was used as a query for BLAST homology search against the “Target syntenic region”, with a minimal coverage of 60% required. When several hits were found, the best hit was kept as a result. When no hit was found, a second BLAST search was conducted against the whole target genome. Proto-genes without hits were annotated.

When a proto-gene got several hits in a target line, only the best hit was considered for further analysis. Final best BLAST hits were referred to as “non-coding homologs”. One query proto-gene can have a maximum six “non-coding homologs”, in a maximum of six outgroups lines.

### Step 3. Assessing mechanisms of proto-gene birth

All “non-coding homologs” were aligned to their proto-gene. For all query proto-genes which contained an intron, the intron was removed in the alignment between the proto-gene and its non-coding homolog, in both sequences. Several diverging features were investigated between the aligned proto-gene and “non-coding homologs”. We investigated the presence of the following six features (SuppData Figure 2): i) start codon, ii) stop codon, iii) frameshift mutation, iv) anticipated stop codon, v) different sequence hit size, and vi) transcription event. When the “non-coding homologs” did not align to the entire sequence of the proto-gene, meaning that the match started later in the sequence and/or ended earlier, the “non-coding homologs” was annotated as “Different sequence hit”. When the size of the alignment was the same, the start codon, stop codon, and frameshift mutations were searched for. The presence of an anticipated stop codon was searched in the sequence. Finally, the genomics positions of the “non-coding homologs” were mapped to their reference genome, their orientation in the genome was assessed, and the presence of a transcription event was annotated.

### Programming and analyses

All statistical analyses were performed with R (Team et al., 2013; Giorgi et al., 2022). Data reshuffling, search and analyses were performed with python (Van Rossum and Drake, 2009), and the following modules were used: Biopython (Cock et al., 2009), matplotlib (3.5.0) (Caswell, 2021), numpy (1.20.3) (Harris et al., 2020), pandas (1.3.3) (Reback et al., 2020), seaborn (0.11.2) (Waskom, 2020), gffutils (0.10.1) (http://daler.github.io/gffutils). All programs and scripts are available in github (available after publication).

### Data access

Our genomic data set is available under NCBI Bioproject accession (Link to be provide, in progress). Additionally, a package containing processed data will be available after final publication, and is referred in the main text as “supplemental deposit”. All programs are stored in the github link (accessible after publication).

## Competing interest statement

The authors declare no competing interests.

## Acknowledgments

We thank Claudia Fricke for her help with fly breeding and maintenance. We are very grateful to the DrosEU consortium, especially Elio Sucena, Josefa Gonzalez, and Iryna Kozeretska for providing European fly lines, and to John Pool for providing the African fly line. We thank Pablo Mirat for constructive feedback on the manuscript. We thank Margaux Aubel for her feedback on the figures.

## Authors contribution

AG and EBB conceptualised the study; JP imbred and prepared the lines, AG and KB extracted the DNA and RNA, AG sequenced the DNA; AG and ML assembled the genomes, AG detected proto-genes, AG and LK studied non-coding homologs, AG, EBB and JP validated and curated the datas, AG and EBB wrote the original draft, AG, EBB, JP, ML, LK and KB corrected the draft, AG and EBB acquired funding acquisition. All authors have read and agreed to the published version of the manuscript

## funding

AG and EBB acknowledge funding by Alexander von Humboldt-Stiftung. This work was supported in part by the Deutsche Forschungsgemeinschaft priority program “The genomic basis of evolutionary innovations” (SPP2349; project number 503272152 awarded to JP (GZ:PA 903/12 -1) and EBB (GZ:BO 2544/20 -1).

## Notes

### Competing Interest Statement

The authors have declared no competing interest.

